# The BET inhibitors JQ1, AZD5153, and I-BET151 co-opt ubiquitin proteasome system components for altering expression of the BRD4 interactome in a human B cell line

**DOI:** 10.1101/2023.11.09.566482

**Authors:** Girish Nallur

## Abstract

Chemoproteomic analysis of the BET inhibitors, JQ1, AZD5153, and I-BET151, identified an extremely large signature of ubiquitin modified proteins associating *in vitro* with a recombinant BRD4 N-terminal protein fragment. The identified proteins included those with known functions in BRD4 complexes for transcriptional and epigenetic control (acetylated histones, the MED complex, BAF complex, RNA pol II transcription complexes, and chromatin-associated complexes). The BRD4 interactome in response to BET inhibitors is suggested to be orchestrated by compound-specific differential actions of up to 16 E3 ligases, 4 deubiquitinase enzymes, and 51 accessory proteins of the ubiquitin proteasome system (UPS). The UPS response of BET inhibition also involves proteins necessary for Myc enhancer binding and Myc response gene expression. A large cohort of UPS substrates commonly responsive to JQ1 and AZD5153 treatments suggests the existence of distinct mechanisms, one involving compound-activated UPS proteins, and another via their direct actions on BRD4. The findings raise the intriguing possibility that UPS triggers promoting proteostasis changes to the BRD4 interactome may be mechanistically coupled with BRD4 function in a proximity-dependent, chromatin-associated manner. Consequently, BET inhibitors and their downstream effects present highly complex environments which may lead to polypharmacology, the phenotypic outcomes or overall clinical benefits of which are hard to assess. However, many new targets and small molecule combinations suggested in this study may afford a path forward for narrowly and more selectively targeting Myc in the clinic with potentially cleaner profiles compared with BET inhibitors or BRD4 as target.

## INTRODUCTION

The transcription factor and growth regulator MYC is essential for normal development but even small increments in MYC expression are potent drivers of many cancers [Zaytseva & Quinn, 2017]. Understanding the multitude of growth signaling pathways and proteomic networks which regulate *MYC* in health and disease will provide new insights and a therapeutic strategy for treating *Myc*-driven malignancies. Numerous studies [reviewed in: Wolf *et al*, 2015] have outlined the tight regulatory controls and transcriptional patterning inherent in fine tuning the functions of Myc and its dimerizing partners.

BRD4 drives MYC transcription by binding to a super-enhancer 1.7 Mb downstream of the *MYC* promoter as well as by recruiting the transcription factor pTEF-b, and associating with chromatin with the help of the mediator complex [MEDs] [Shi *et al*, 2013]. The repeated failures of bromodomain and extraterminal (BET) inhibitors in the clinic, including JQ1 whose structure mimics acetylated lysine, suggests that many additional biological factors may be involved which account for as yet unknown phenotypes and drug resistance. Approaches for finding the mechanisms and proteins involved in BET inhibitor functions may help develop more effective clinical regimens [Shorstova *et al*, 2021, Filippakopoulos *et al*, 2010]. Many additional BET inhibitors and PROTACs derived from BET inhibitors are being studied in the clinic.

While roles of BET inhibitors at the transcriptional and chromosome maintenance levels are fairly well understood, relatively little is known if these compounds directly or indirectly impact protein homeostasis via the UPS. Commonly observed proteomic changes include synthesis, maturation, sub-cellular localization, and intrinsic degradation of proteins or their network partners [Juskiewicz & Hegde, 2019]. Such proteins functioning in the mechanistic paths of BET inhibitors can potentially impact BET inhibitor outcomes in the clinic and characterizing them can help better understand the complexity. This additional layer of information is not only useful but imperative for the design of PROTACs and small molecule compositions for targeting Myc [Bekes *et al,* 2022]. Further, proximity-mediated, mechanism-based, and compartment-specific UPS events may be contributing factors. These micro-events may be hard to detect or quantitate over the larger cellular pools of the same proteins, particularly when the same protein is stable elsewhere. The ability to observe these effects at a single protein level over space and time and integrating such information with appropriate chemistries will help build a pipeline of mechanistic targets and compounds for more effective therapies.

The UBQuest proteomics platform addresses the above factors at the whole proteome level then deconvolutes the information to the single protein levels with mass spectrometry. It is a proximity-based, preparative process for forming, enriching, separating, and purifying complexes of E3 ligases (E3Ls) along with their engaged substrate(s), with simultaneous elimination of all other proteins in the sample. The product of UBQuest is a pool of proteins comprising multiple E3 ligases, the engaged substrates, deubiquitinases, proteasome subunits, substrate recognition adaptor proteins, ubiquitin-like proteins, SUMOylated, neddylated, and ubiquitinated proteins, ubiquitin-binding proteins, and any other proteins participating in substrate engagement (the active ubiqitome). Quiescent E3Ls and deubiquitinating enzymes (DUBs), UPS substrates, or any proteins not involved in substrate interaction are eliminated during the UBQuest process, regardless of their co-expression. The highly enriched pool of proteins are characterized with mass spectrometry, and is a source for co-precipitating interacting partners.

In this initial report, chemoproteomic analysis of three BET inhibitors in a human B cell line is presented. In this study, it is understood that the 3 inhibitors employed (JQ1, AZD5153, and I-BET151) exhibit different modes of action, potency and specificity (bivalent vs monovalent, selectivity for different BRD containing proteins, etc.). In addition, the DOHH2 cell line employed may impact the biology of Myc differently than some other cell lines. So, a full basis for comparing the proteomic signatures for drug discovery applications requires finding an equivalence point to be clinically significant.

However, the focus of this study was to utilize the BET inhibition strategy as a means of altering the physiology of cells, and the protein networks therein, including the BRD4 physical interactome. This approach provided samples for implementing the assays in the present study. It follows that the same process can be iterated with well-defined clinical samples for observing protein networks *per se* or in comparative mode using multiple compounds. Therefore, the associations presented herein must be viewed as defining the assay characteristics, rather than for their clinical significance.

Several components of the UPS were involved in the response to BET inhibition, as determined with mass spectrometry. Proteins complexing with BRD4 were highly enriched, suggesting *in vitro* complex formation with UPS modified proteins. JQ1, AZD515 and I-BET151 each exhibited their own proteomic signatures and E3 ligase specificities suggesting different mechanistic paths and phenotypic outcomes. The common footprint of JQ1 and AZD5153 action was most notable, suggesting downstream effects of BRD4 on its own interactome likely via a combination of kinase activity and ubiquitination. Overall, the multitude of proteomic alterations observed with transcription and epigenetic factors present a highly complex molecular signature which may impact BRD4 function and Myc responses. The potential applications of the process and some findings relating to BET inhibition in the clinic are discussed.

## RESULTS

### Chemoproteomic analysis identifies signatures of E3 ligases and their putative substrates in BET inhibited cells which associate *in vitro* with unphosphorylated BRD4 N-terminal fragment

The human diffuse large B cell lymphoma (DLBCL) cell line DOHH2 was treated with the BET inhibitors JQ1, AZD5153, or I-BET151. Each drug-treated sample was subjected to the UBQuest assay and BRD4-associating UBQuest proteins were purified by *in vitro* association with a bacterial recombinant protein fragment of BRD4 containing the BD-I and BD-II domains (BRD4-N, 49-460aa). An outline of the UBQuest-BRD4-N selection process is shown in Fig. 1. The resulting, doubly selected proteins characterized by E3 ligase action and BRD4-N physical association, termed UBQint, were identified with mass spectrometry.

**Fig. 1:**
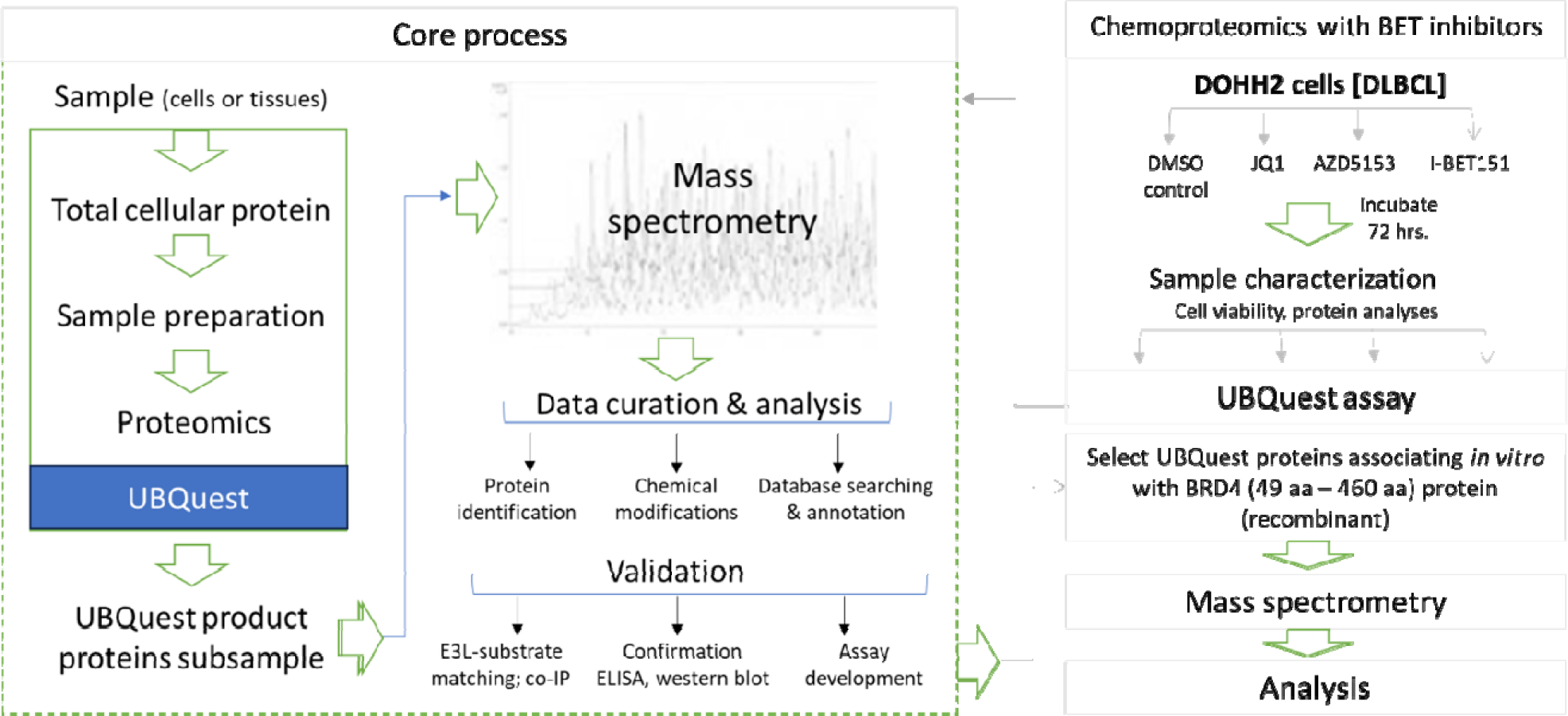
The box marked ‘Core process’ shows the general workflow and key steps in the UBQuest process. The box on the right shows the workflow for iterating UBQuest with BET inhibitor treated DOHH2 cells.

A combined 1,286 unique proteins were identified by mass spectrometry across the 4 UBQint samples, represented by 25,899 analyzed peptides (Table I). Other than proteins with UPS or endoplasmic reticulum-associated degradation (ERAD) functions (see below), about 15-18% of the UBQint proteins in the DMSO, JQ1 or AZD5153 treated samples are known from the literature to complex with BRD4. The remaining proteins may be part of larger protein complexes formed during the process involving chaperone proteins, networking, or ubiquitin- or SUMO-binding proteins. Only 10% of the proteins in the I-BET151 sample were BRD4 interacting, suggesting decreased ubiquitination activity, BRD4 phosphorylation or dimerization promoted by I-BET 151 compared with the other BET inhibitors.

**Table I.** Summary information from the mass spectrometric analysis of UBQint proteins with BET inhibition. Column 2: total number of peptides analyzed from each of the samples shown in column 1, and the number of unique proteins they represent (column 3).

| Compound | Number of peptides analyzed | Number of unique proteins in sample | Number of proteins known to complex with BRD4 [from literature] |
| --- | --- | --- | --- |
| DMSO [control] | 4892 | 494 | 84 |
| JQ1 | 10212 | 1138 | 162 |
| AZD5153 | 9580 | 1100 | 163 |
| IBET-151 | 2234 | 455 | 76 |
| **TOTAL** | **26918** | **1340** | **194** |

### UBQuest detects compound-induced changes to UPS system components and their putative substrates

Analysis of the distribution of UBQint proteins identified 112 proteins unique to JQ1 treatment, 127 proteins with AZD5153 treatment, and 17 proteins with I-BET151 treatment (Fig. 2a). Distinct UPS proteins were present in each subsample, suggesting unique mechanisms of UPS responses triggered by them. Preliminary analyses of the proteins allowed them to be grouped into proteins sharing common amino acid signatures (‘motifs’) in their primary sequences. It remains to be determined if the motifs represent E3 ligase recognition sites or the sites recognized by enzymes effecting other types of post-translational modifications. It is also not clear if these are off-targets of the corresponding compounds, but the identification of motifs suggests a common origin. 24 proteins identified uniquely in the DMSO subsample were housekeeping proteins, with the exception of 2 transcription factors. As previously noted, the DMSO subsample did not contain any UPS proteins, but included 9 proteins known to interact with BRD4. At 37.5%, BRD4-interacting proteins are highly enriched in the DMSO subsample compared with the average across 4 samples. It remains to be seen if loss of these proteins is a contributing factor in BET inhibition. The 266 proteins common to all samples were housekeeping proteins.

**Fig. 2a.**
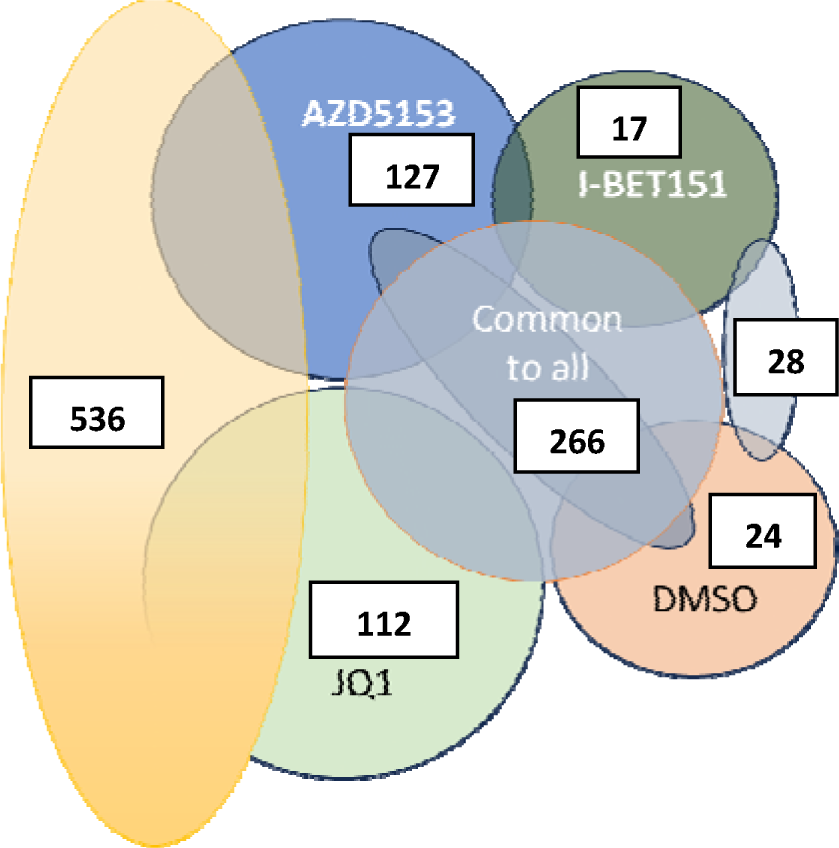
Venn diagrams showing the combined or individual responses of BET inhibitors on UBQint proteins. The 536 proteins common to JQ1 and AZD5153 treatments was the single largest set, followed by the set of 266 proteins which remain unchanged with BET inhibition. These were mostly housekeeping proteins.

The UBQint proteins were annotated with their functions suggested in the literature. As shown by the grey bars in Fig. 2b, 69 UBQint proteins are components of the UPS. Of these, 18 proteins are E3 ligases, 4 are DUBs, 21 are subst rate r ecognition adaptors, chaperones, or proteins, 5 are E3 ligase modifying proteins, 2 are ubiquitin-like proteins, 9 are ubiquitin binding proteins, 8 are proteasome subunits, 3 are SUMO proteins, and the last is ubiquitin. Ubiquitin comprised the largest single protein in this group represented by 230 peptides, suggesting that many UBQint proteins were tagged with it. The UPS group of proteins comprised 1,989 peptides representing 8% of the total peptides in the study. The large enrichments of UPS modified proteins in UBQint samples follows the purpose for which the UBQuest process is designed - co-purification of UPS proteins and their engaged substrates directly from biological samples.

**Fig. 2b.**
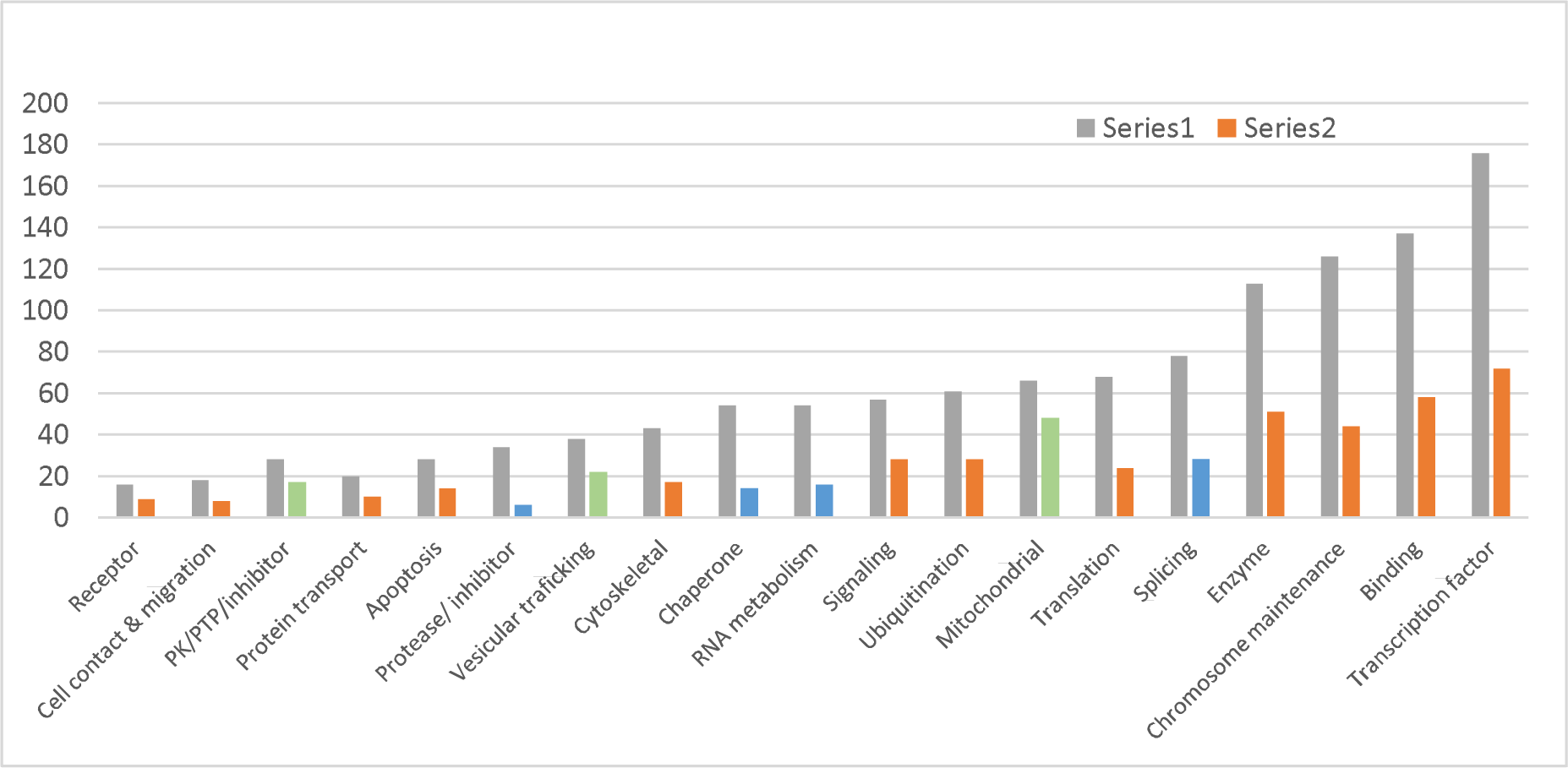
Representation of known functions of UBQint proteins. Number of proteins of a specific function identified across all 4 samples (grey bars); commonly and uniquely in JQ1 and AZD5153 (orange bars); over-represented in JQ1 and AZD5153 compared with the average for all 4 samples (green bars); and under-represented in JQ1 and AZD5153 compared with the average for all 4 samples (blue bars).

A solitary E3 ligase and a DUB were commonly identified in all 4 samples and they were the only E3L and DUB in the DMSO-treated sample. In sharp contrast, 10 E3 ligases and 2 DUBs were enriched commonly in JQ1 and AZD5153 treated samples and the number of putative substrates identified in these samples was correspondingly larger (see below). Five E3 ligases and 1 DUB and 1 E3 ligase were unique to JQ1 the AZD5153 treated samples, respectively in addition to other UPS proteins. These observations suggest that BET inhibitors may collaborate with components of the UPS and ERAD, which explains their enrichment with UBQuest along with co-purification of the substrates (see below).

UPS proteins were under-represented in the I-BET sample, with only 107 peptides (5% of the total UPS peptides in the study) representing 23 UPS proteins. These included 3 E3 ligases, 8 E3L adaptors, 3 ubiquitin binding proteins, 4 proteasome subunits, 3 ubiquitin binding proteins, and 2 ubiquitin-like proteins. Overall, I-BET151 peptides and proteins resembled DMSO, suggesting that the compound did not significantly alter cellular UPS activity, or diminished assay robustness compared with JQ1 or AZD5153. Perhaps, utilizing the full length BRD4 protein, phosphorylated BRD4, or dimerized BRD4 in the selection step would detect more robust UPS signatures (see Fig. 8). Nevertheless, these data reaffirm the utility of UBQuest for selectively purifying proteins modified by UPS activity and as a tool for characterizing compound interactions with UPS system components in biological samples.

Strikingly, a small but significant number of BRD4 peptides identified in the sample extend beyond the 460aa terminus of the recombinant BRD4-N protein fragment, showing that native BRD4 protein was recruited into the complexes via dimerization or complex formation. Some UBQint proteins were likely enriched by complexing with the BET and extraterminal domains through to the C-terminus of native BRD4 (see below).

While a comparable number of proteins were identified in control experiments involving direct BRD4-N *in vitro* selection and elimination of the UBQuest step, these proteins showed poor correlations with UBQint proteins. Native BRD4 protein was not detected in the control samples and substantially fewer had known interactions with BRD4. These observations suggest that protein pools participating in the control complexes are distinct from those in UBQint, perhaps arising from uncontrolled protein interactions rather than being driven by proteins possessing unique modifications, as in the UBQint proteins (data not shown). These observations suggested that UBQint proteins may be a distinct subset of the proteome in BET inhibited cells whose protein modifications served as a bridge or driver for effective protein complex formation *in vitro*. The fidelity of such interactions or if the same interactions occur in cells remains to be confirmed.

### Identification of acetylated histones and proteins affecting histone metabolism and nucleosomal recruitment in UBQint proteins

BET inhibitors mimic acetylated histones and a suggested role for their mode of action is that they interfere with BRD4 binding to acetylated histones associated with chromatin [Wernersson *et al*, 2022]. Therefore, UBQint proteins were queried for the presence of proteins belonging to this group.

Seven histones (3 acetylated), 9 enzymes involved in histone acetylation or deacetylation, and alterations of the methylation status of chromatin, and 10 proteins which influence the organization of histones in nucleosomes were identified, within a total signature of 91 proteins. The set also included 6 DNA polymerase subunits, 22 proteins involved in DNA replication and repair, 13 proteins involved in mitosis, and 7 proteins involved in checkpoint control of chromatin remodeling. Curiously, any of these proteins differed in their response to BET inhibitors based on their peptide sequence signatures from mass spectrometry. The reasons for the differences and their significance to BET inhibitor actions or cell type specificity need to be understood with more experimentation.

### JQ1 and AZD5153 possess common features of inducing changes to UPS system components and their putative substrates

Additional clues that BET inhibitors may promote changes to E3 ligase activity were suggested by observing the distribution of UBQint proteins in different subsamples of this study. Nearly half of the proteins across the 4 samples were specific and common between JQ1- and AZD5153-treated subsamples (termed JQ1-AZD; 536 unique proteins; Fig. 2b, series 2). While the putative functions of JQ1-AZD proteins followed the same pattern as those for the entire sample, (Fig. 2a, series 1, gray bars), mitochondrial proteins, protein kinases/phosphatases, and vesicular transport proteins were over-represented (Fig. 2a, green bars), while RNA binding proteins, chaperones, and protease inhibitory proteins (green bars) were under-represented. Myc-response gene products were under-represented – there were only 10 out of 43 total in the 4 samples (details below).

Importantly, on average the number of optimized peptides (see below for definition) per protein in the JQ1-AZD subsample was only 7.4 (total of 3,974 peptides in subsample), while only 5.7% of these proteins are known to interact with BRD4, compared with the sample average of 20 peptides and 14.5%, respectively, for the entire sample. These findings suggested that the UPS response was significantly higher with JQ1 and AZD5153 treatments over other samples and consistent with the larger number of UPS proteins identified in this subsample. Consequently, substantially more proteins were likely to be enriched with the JQ1-AZD subsample and ubiquitin modification increased the affinity of complexing and networked interactions in the BRD4-N selection step.

BRD4 been shown to possess intrinsic protein kinase activity [Weissman *et al*. 2021] which resides in three of its domains, including BD-II, and maps to amino acids 351 to 598. BRD4 phosphorylates the C-terminal domain of RNA polymerase II, Myc, TAF7, NSD3, CDK9, and MMLV virus integrase proteins. Further, Deviah *et al*. [2012] showed that BRD4 phosphorylation of Myc at Ser58 promotes its degradation, suggesting phosphorylation-triggered, ubiquitin-mediated degradation as a potential mechanism of BRD4 kinase. Myc was not detected in the UBQint samples; however, the set of 49 proteins mapping to the BRD4 zone (Fig. 2d.) included TAF7 and 3 protein kinases other than BRD4. Proteomic assays and peptide analyses confirmed adjacent phosphorylation and ubiquitination sites in some of the 49 proteins (data not shown), suggesting phosphorylation and ubiquitination to be linked, concurrent, or proximity-dependent events with E3L-substrate co-localization being a prerequisite. These observations also point to the likelihood that BRD4 kinase activity may be involved in phosphorylation of some of the JQ1-AZD proteins. Other types of protein modifications are suggested in some proteins in this set with reference to the PhosphoSitePlus database (version 6.7.1.1) including, acetylation, methylation, SUMoylation, and monoubiquitination. The significance of these modifications to BRD4 function needs to be elucidated.

To address the question of compound- or BRD4-mediated changes accounting for JQ1-AZD proteins, the associated data were analyzed with reference to a hypothesis to allow construction of a model for testing and reconfirmation. Assuming that the biological factors influencing the abundance of the proteins (including protein resynthesis rates) and residual biological activity of BRD4 in the presence of JQ1 and AZD5153 are the same, the optimized peptide count for any protein in this subsample is expected to be identical if UPS activity is influenced by BRD4. However, the peptide count can be different if ubiquitination is compound-specific *via* interactions with UPS components or substrates, independently of the compound’s action on BRD4.

For the above purpose, peptide signatures of the JQ1-AZD proteins from mass spectrometry were optimized. Peptides with identical sequence for each protein were quantitatively subtracted between the JQ1 and AZD samples, and the remaining unique peptides were termed optimized peptides (henceforth termed ‘peptide counts’). This process eliminated peptides arising from common tryptic signatures without E3 ligase actions as well as any E3 ligase(s) which may have acted commonly on the same protein in the two subsamples for routine turnover, but retained peptides whose effects on the protein were likely specific to JQ1 or AZD5153 treatments, or both.

The peptide count for each of the 536 JQ1-AZD proteins in the AZD5153 sample were plotted against the count for the same protein in the JQ1 sample and expressed as a percentage ratio (JQ1/AZD). As shown in Fig. 2c, the ratio plot yielded a chair-shaped line with 2 inflection points the first of which involved 174 proteins on the left while the second on the right involved 226 proteins, with a flat line (peptide percent ratio of 100) in the middle involving 206 proteins.

**Fig. 2c.**
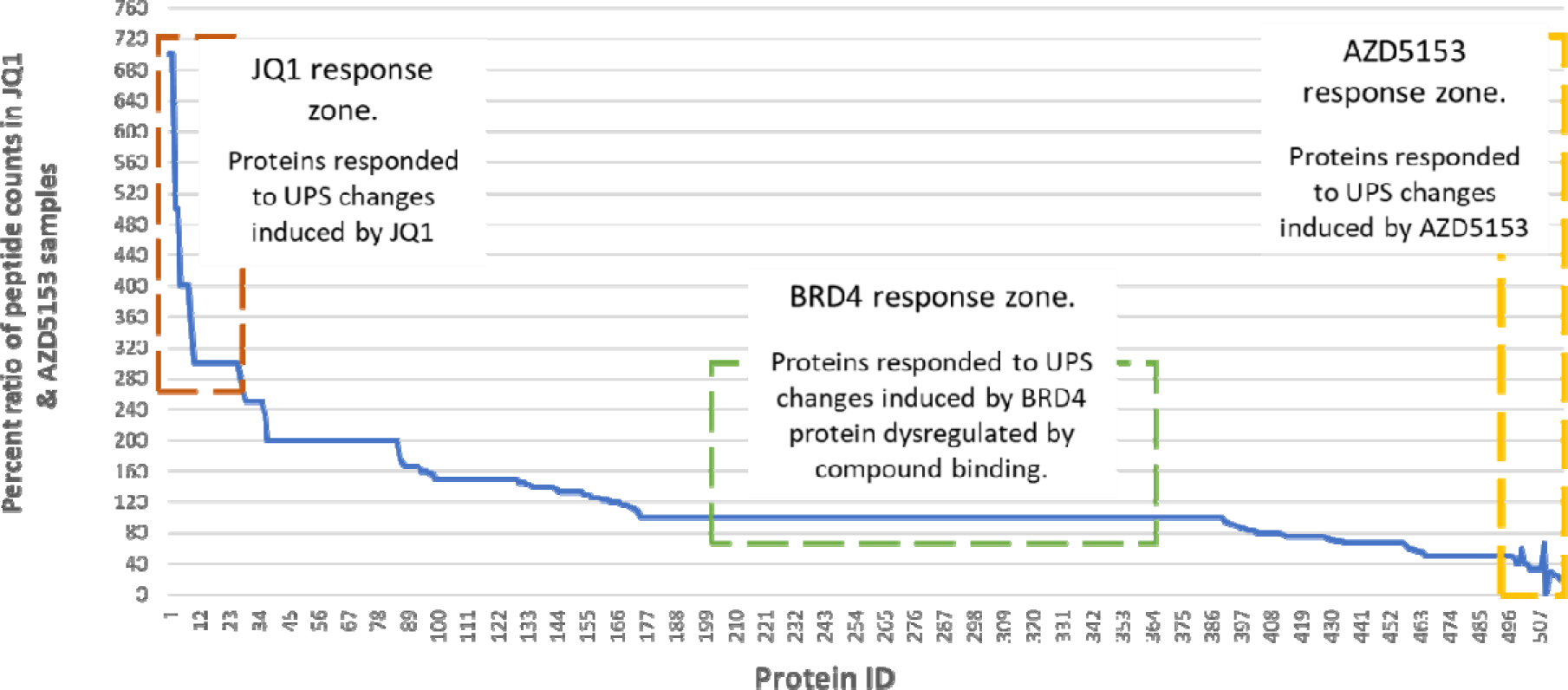
A chart showing how UBQint proteins common to JQ1 and AZD5153 treatments were segregated into different response zones using the prediction model. The X-axis represents protein candidates, Y-axis shows the per cent ratios of optimized peptide counts for each protein. Proteins mapping to the orange box, yellow box, and green box indicate protein candidates responding to JQ1, AZD5153, and BRD4, respectively.

**Fig. 2d.**
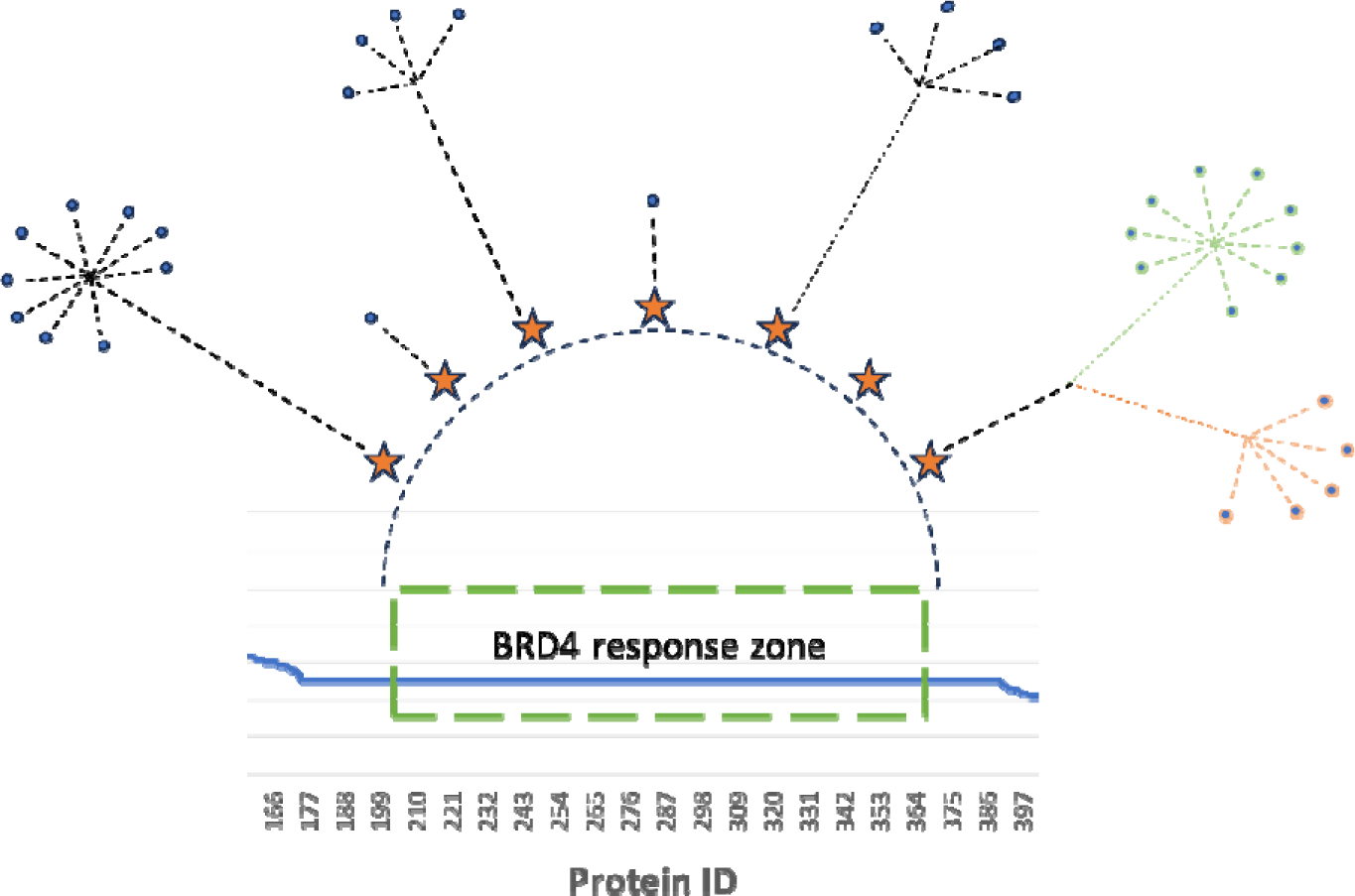
Annotation and first level interactions of the 49 proteins shown in the BRD4 response zone in Fig. 2c as suggested from literature. Orange stars represent BRD4 interacting proteins. Grey dots indicate first level interactors of the protein represented by the orange star.

Thirty-five proteins were identified from the ratio plot having peptide ratios of 2.5-fold or higher in the JQ1 sample compared with the AZD5153 sample (located at the far left of the ratio plot). These proteins are suggested to be maximally responsive to JQ1 treatment and acting *via* the UPS response, but likely much less with AZD5153. Proteins affecting the tumor microenvironment and plasma membrane structure, 8 proteins with mitochondrial functions, and 8 proteins with nuclear functions were identified in this group. Many of these proteins are implicated in AML, myelodysplastic syndromes, hematopoietic stem cell maturation, autoimmune diseases, inflammation, and viral infections.

Fourteen proteins with functions in the nucleus are suggested to be maximally responsive to AZD5153 treatment (located at the far right of the ratio plot). Proteins involved with the BAF complex, nuclear phase separation, neurodegenerative disorders, tumor suppression, and hematopoietic stem cell differentiation are included in this set. Together, these 49 proteins are provisionally designated as BET inhibitor-responsive proteins whose biology is likely affected in a compound-specific manner by the compounds interacting with the UPS system independently of their actions on BRD4, the significance of which or its validity in cells needs to be confirmed.

To further test the assumptions and preliminary conclusions about BRD4 or compound-sensitivities affecting the JQ1-AZD proteins, a set of 49 proteins mapping to the horizontal portion of the ratio plot (the putative BRD4 response zone) were analyzed. Seven proteins in this set are known from the literature to complex with the BRD4 protein (Fig. 2d). Curiously, 33 of the remaining proteins were also found to be first level interacting partners of the BRD4 interacting proteins, indicating *in vitro* protein complex formation with BRD4 during UBQuest as well as interaction selection with BRD4-N protein. Importantly, the same protein complexes were not observed with untreated DMSO controls or I-BET151 treated cells, indicating a driver effect for protein complex formation and suggesting convergence of the JQ1 and AZD5153 pathways in determining the physiological state of BRD4 as well as the proteins in its interaction network.

Sample analysis of optimized peptide sequences for proteins in the JQ1 and AZD5153 samples showed compound-specific amino acid sequences in the corresponding subsamples, but were identical in proteins in the BRD4 response set, providing internal validation (data not shown). The JQ1-, AZD- and BRD4-response proteins can be profiled separately and validated in cell-based assays and the E3 ligases responsible for each can be co-purified from the appropriate subsamples. If indeed some BRD4 responsive proteins are the substrates of BRD4 kinase, it may help to more closely dissect the mechanisms of action of BET inhibitors to understand phenotypic complexity at the molecular levels. BRD4 kinase targets can also be explored as standalone therapies to eliminate the multiple outcomes of BET inhibition, as suggested by this investigation.

### BET inhibitors alter the proteostasis of transcription factors which collaborate with BRD4

As a transcription factor and epigenetic regulator, BRD4 interacts with several proteins for each function. By binding to acetylated histones associated with chromatin, BRD4 accumulates at lineage specific transcriptionally active regions along promoters and super enhancers (SEs), and promotes transcription by RNA polymerase II at the initiation and elongation stages, particularly in controlling the transcription of Myc. The non-transcriptional roles of BRD4 include DNA damage repair, checkpoint activation, and telomere homeostasis. BRD4 serves as a nucleation center for the assembly of large protein complexes that promote RNA polymerase II activity (Donati *et al*., 2018) involving the Mediator complex, RNA-Pol II subunits, basal and activated transcription factors, and transcription elongation factors.

The earlier presented observations that UBQuest enriches for protein complexes formed *in vitro* prompted an analysis of which of BRD4’s interactors in transcriptional control may have co-purified in the UBQint samples. The Mediator complex is a 30-protein complex necessary for the initiation of transcription. Although many studies have documented protein contacts between the mediator complex and BRD4 [Allen *et al*., 2015; Flanagan *et al*., 1991; Kelleher *et al*., 1990], the exact nature of these interactions are not entirely clear since at least one study found that BRD4 did not co-purify with other Med proteins using chromosome immunoprecipitations (CHIP; Quevedo *et al*., 2019). Resolving this aspect of BRD4 function is critical for clinical applications. While inhibition of Mediator is one of the central mechanisms underlying BETi cytotoxicity, its relevance to the therapeutic effects of BET inhibitors in cancer is unclear [Bhagwat *et al*., 2016]. Besides, the Mediator complex is capable of forming phase condensates essential for super-enhancer functions not dependent on high levels of BRD4 and Mediator, and may include additional activator complexes at some sites, such as, CTCF and cohesin [Crump *et al*., 2021].

To try to resolve the BRD4-Mediator complex interactions question, UBQint proteins were queried for Med proteins which may have co-purified by association with the BRD4-N recombinant protein fragment. Eight members of the Med complex were identified (Table III), with different peptide counts in treatments with JQ1, AZD5153, or I-BET151 (Fig. 3b). Med1 and Med4 were identified in all samples, including the DMSO control sample, but none of the other Med proteins were present in either I-BET151 or DMSO samples. This finding is consistent with the report from Gibbons *et al*. [2019] that treatment of TH1 polarized PBMC cultures with JQ1 did not significantly alter the expression of Med1. The effects of BET inhibitor on expression of other Med subunits in Fig. 3b is less clear. Some of these have been reported to be involved in phase separation of epithelial-mesenchymal transitions in cancer, but rigorous confirmation is lacking.

**Fig. 3a.**
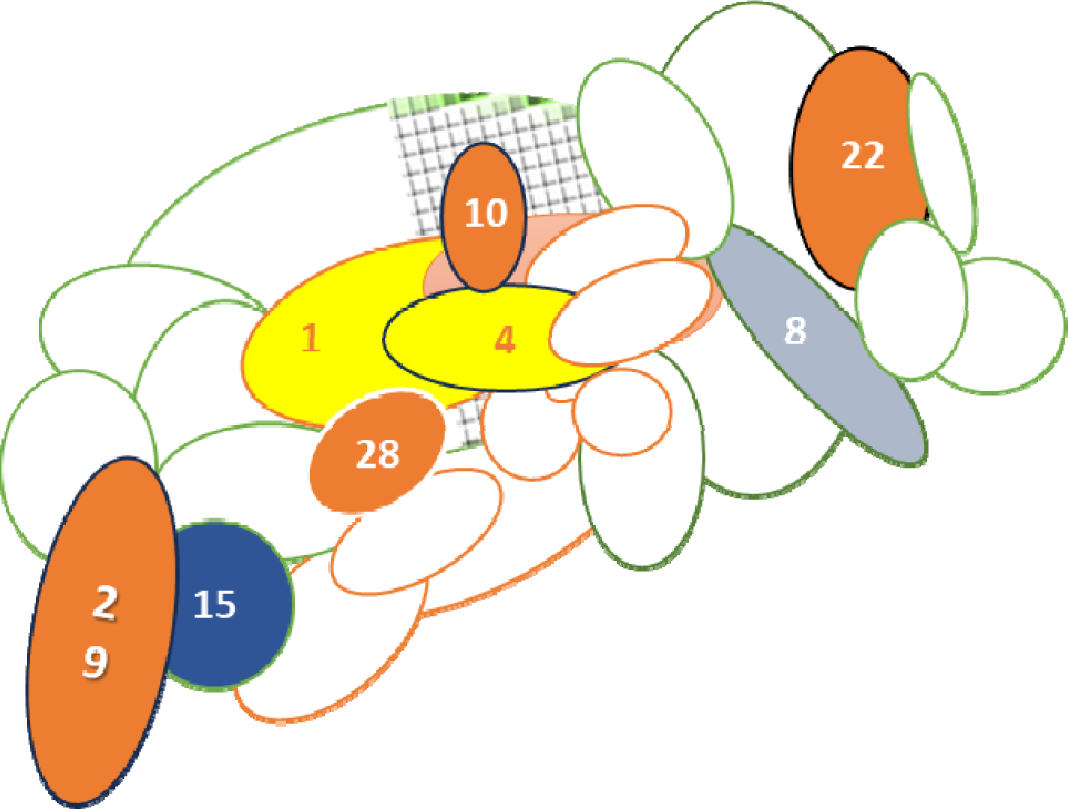
Diagrammatic representation of the Mediator complex proteins identified with UBQint process. Numbers indicate the corresponding MED subunit. Yellow ovals – common to all 3 BET inhibited and DMSO samples. Orange ovals – identified commonly in JQ1 and AZD5153 treated samples. Grey oval – specific to AZD5153, and blue circle – specific to JQ1 treatment.

**Table II.**
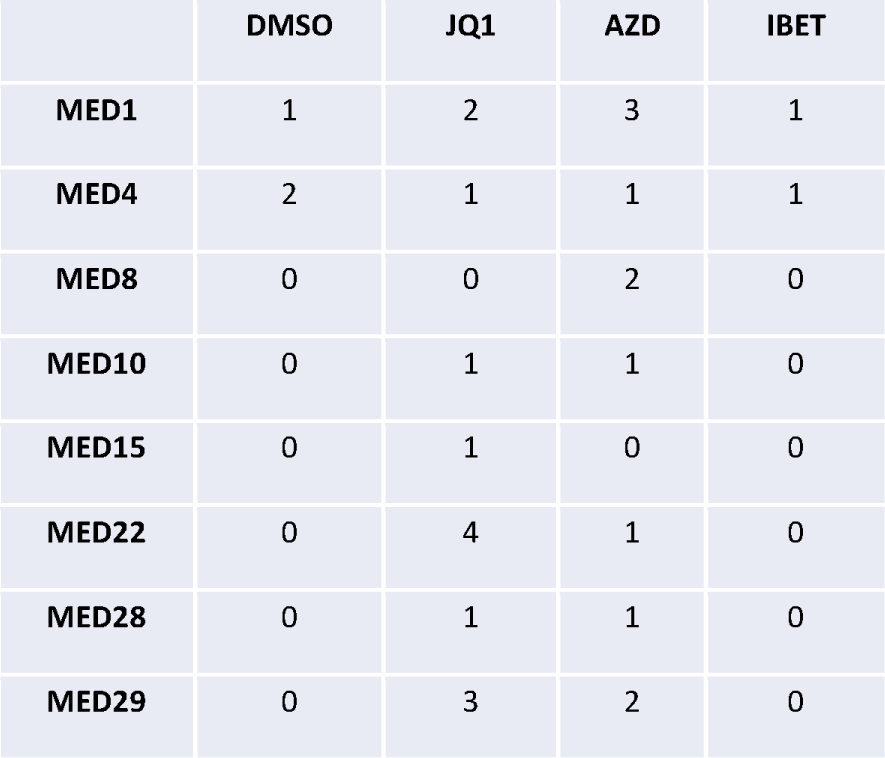
Shows the number of optimized peptides identified for each Med protein in the UBQint samples.

|  | DMSO | JQ1 | AZD | IBET |
| --- | --- | --- | --- | --- |
| MED1 | 1 | 2 | 3 | 1 |
| MED4 | 2 | 1 | 1 | 1 |
| MED8 | 0 | 0 | 2 | 0 |
| MED10 | 0 | 1 | 1 | 0 |
| MED15 | 0 | 1 | 0 | 0 |
| MED22 | 0 | 4 | 1 | 0 |
| MED28 | 0 | 1 | 1 | 0 |
| MED29 | 0 | 3 | 2 | 0 |

**Table III.** A listing of the UBQint proteins shown in Fig. 4. Column 2 shows the suggestion from literature regarding interaction with BRD4. Columns 3 through 6 show the representation of optimized peptides for each protein shown in column 1 in presence of BET inhibitor (shown in row 1).

| **Protein** | **BRD4 interaction** | **DMSO** | **JQ1** | **AZD** | **IBET** |
| --- | --- | --- | --- | --- | --- |
| GTF2F1 | No | 11 | 17 | 19 | 4 |
| GTF2F2 | No | 2 | 4 | 3 | 2 |
| GTF2I | No | 0 | 2 | 3 | 0 |
| POLR2D | Yes | 0 | 4 | 2 | 0 |
| POLR2F | No | 0 | 1 | 1 | 0 |
| POLR2I | No | 0 | 3 | 2 | 0 |
| POLR2J3 | No | 0 | 0 | 1 | 0 |
| POLR2M | No | 0 | 6 | 2 | 0 |
| PAF1 | No | 0 | 0 | 1 | 0 |
| RTF1 | Yes | 2 | 0 | 0 | 0 |
| LEO1 | Yes | 5 | 6 | 5 | 1 |
| HTATSF1 | No | 0 | 1 | 1 | 0 |
| NELFA | No | 0 | 4 | 3 | 0 |
| NELFE | No | 5 | 9 | 4 | 2 |
| SUPT5H | Yes | 0 | 2 | 2 | 0 |
| CSTF2 | No | 0 | 15 | 21 | 0 |
| CCNT1 | Yes | 0 | 1 | 0 | 0 |
| HEXIM1 | Yes | 1 | 4 | 3 | 1 |
| IWS1 | Yes | 0 | 1 | 1 | 0 |
| SSRP1 | Yes | 1 | 0 | 0 | 0 |
| TCEA1 | Yes | 13 | 15 | 12 | 7 |
| TCERG1 | Yes | 0 | 4 | 4 | 0 |
| UBAP2 | No | 4 | 29 | 20 | 12 |
| UBAP2L | No | 37 | 49 | 41 | 28 |

The most notable observation from the data in Fig. 3b is that Med1 and Med4 may be minimally sensitive to BET inhibition, but that other subunits may be differentially modulated by JQ1 and AZD5153. The peptide counts likely represent stoichiometries in Med protein complexes rather than the absolute amounts of proteins in the BET inhibited samples. The lack of peptides for Med8, 10, 22, 28, and 29 in the I-BET151 and DMSO samples are likely all-or-none effects and not a dose response issue. At least 2 subunits of the cohesin complex were also detected in the JQ1 and AZD5153 samples, located in the JQ1 zone (not shown). Preliminary follow-on data indicates potential contributing factors which modulate turnover of Med proteins, including BET inhibitor-induced E3 ligase switch, activation of new E3 ligases, signaling from the extracellular milieu, or other complexing proteins (data not shown). These observations need to be tested under appropriate physiological conditions and confirmed by purifying the E3 ligase.

Further analysis of the UBQint data identified 24 proteins known to form complexes at promoters containing BRD4. As shown in Table III, these include 5 POLR II subunits, 3 members of the TFIIF basal transcription factor complex, 3 members of the PAF1C complex (PAF1, LEO1, and RTF1) which has multiple functions during transcription and pluripotency of embryonic stem cells, TCEA1 and HTATSF1, which collaborate with the PAF1C complex for transcription elongation, SUPT5H and two subunits of the NELF complex, NELFA and NELFE, which promote transcriptional pausing, CCNT1, a subunit of P-TEFb, and HEXIM1 which regulates P-TEFb to cause transcriptional pausing, TCERG1, a repressor of transcriptional elongation, and 2 proteins which recruit transcription factors to the RNA Pol II complex, UBAP2 and UBAP2L. IWS1, which defines the composition of the RNA polymerase II (RNAPII) elongation complex, and SSRP1, a member of the FACT transcription elongation complex were also detected. Surprisingly, TFIIS subunits were not detected in UBQint proteins. While several isoforms of histone H2B were identified, neither RNF20/40, reported as the E3 ligase which recruits PAF1C for H2B monoubiquitination, nor UBE2A/B were present in the UBQint proteins, suggesting the existence of alternate mechanisms. The impact of BET inhibitors on these proteins, the multiple molecular phenotypes arising therefrom, and the resulting complexes need to be further understood.

**Fig. 4.**
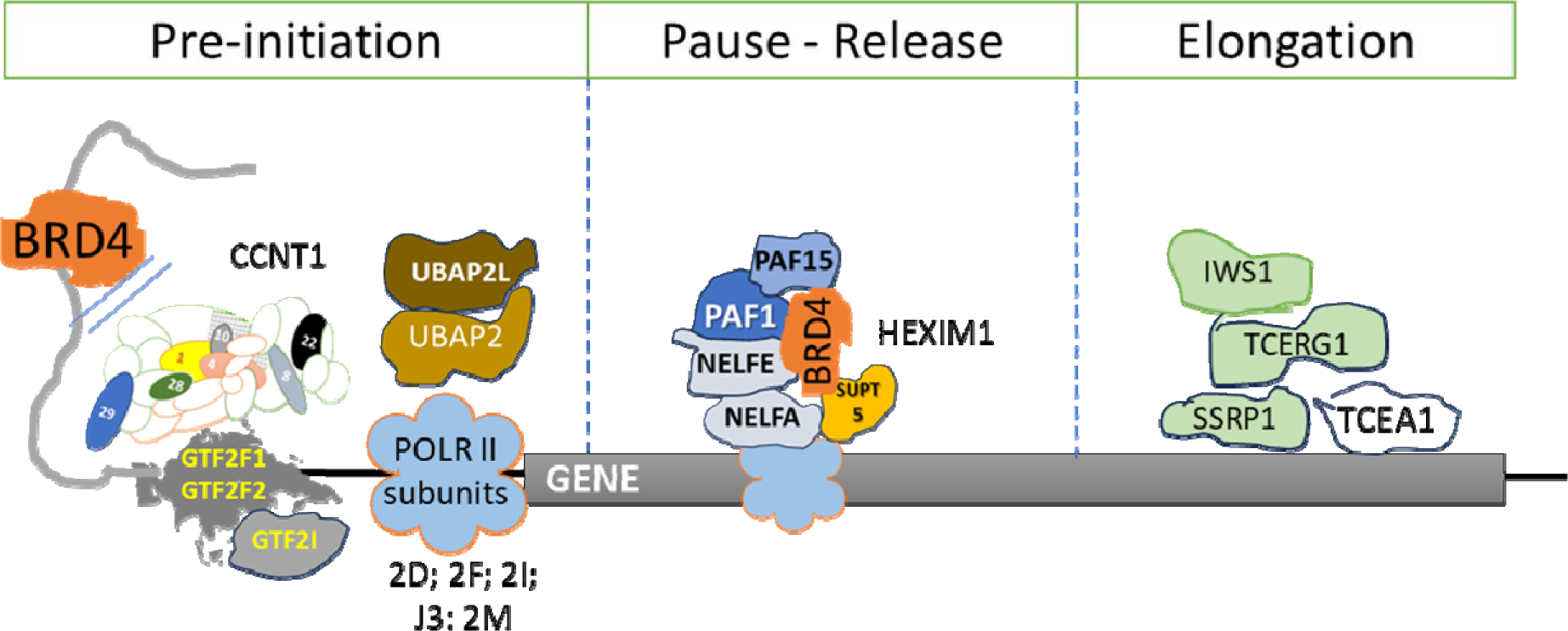
Proteins known in literature to be associated with BRD4 for transcriptional regulation. The diagram shows the proteins identified with UBQint process which are involved in various stages of transcription.

The GTF2F complex proteins GTF2F1, GTF2F2, and GTF2FI appear to be the most impacted by UPS changes in the BET-inhibited samples. GTF2I coordinates transcriptional events at multiple promoters in response to serum stimulation, so it could be an important point of entry for BET inhibitor action. This protein maps to the mid-point of the proposed JQ1 and BRD4 response zones on the ratio plot shown in Fig. 2c., so it appears that its regulation may be complex. Despite comparable peptide counts for GTF2F1, preliminary data suggest major differences in UPS targeting of this protein by each of the BET inhibitors (data not shown).

Many additional differences exist as shown in Table III; however, the increased association with UBAP2 is most notable since it, along with UBAP2L, recruits other proteins to RNA pol II and may be chiefly responsible for building chromatin-associated multiprotein complexes. A few transcriptional repressors are also present among the proteins in Table III, the differential proteostasis of which can appreciably alter the phenotypes of BET-inhibited cells. Profiling these proteins in combination with RNAseq can help determine which genes are affected.

In an excellent study, Sabari *et al*. [2018] reported that the intrinsically disordered regions of BRD4 and Med1 form condensates at super enhancers, resulting in liquid-liquid phase separations and help form clusters which are in close spatial proximity. Many additional proteins and enzymes necessary for the formation and regulation of phase separation, and the involvement of proteins rich in proline and glutamine in this process have been reported in the literature, many of which were also present in the UBQint samples. Using structural studies, Babu *et al*. [2022] identified that the enzyme peptidylprolyl isomerase A (PPIA) concentrates inside liquid-like droplets formed by the Alzheimer’s disease-associated protein tau, as well as inside RNA-induced coacervates of a proline-arginine dipeptide repeat protein, PR20. Curiously, PPIA represented by 286 peptides, was one of the most highly enriched proteins with the UBQint process. The significance of these findings and their impact on transcription functions of BRD4 will help better characterize the mode of action of BET inhibitors.

### BET inhibitors influence differential proteostasis of wild type forms of the FET fusion oncoproteins, FUS, EWSR1, and TAF15

FET oncoproteins are formed as a result of chromosomal translocations juxtaposing them with transcription factors resulting in fusion proteins which are pathogenic in sarcoma and leukaemia. They co-localize with BRD4 on chromatin*via* direct binding and shared interactions with the SWI/SNF complex [Linden *et al*., 2022] affecting chromatin structure and phase separations. While the FET fusion oncoproteins affect mechanisms of action of many drug candidates, including BET inhibitors and the PROTAC ARV-825 [Chen *et al*., 2019], their impact on the wild-type forms of FET proteins is relatively unknown.

Wild type FUS and EWSR1 proteins appear to interact in a similar manner with BET compounds based on peptide counts in the UBQint samples but compound-specific differences in sensitivities to E3 ligases are likely. TAF15 mapped in the interval between the BRD4 and AZD5153 response zones on the ratio plot shown in Fig. 2c., suggesting influences from BRD4 as well as AZD5153 for protein modification affecting either its stability or interaction with a complex containing BRD4, and consistent with the higher peptide counts in that sample. Like FUS and EWSR1, TAF15 binds to and effects the turnover of more than 4,900 RNAs [Kapeli *et al*., 2016]. The authors noted that shRNA mediated targeting of TAF15 in inhibitor-treated iPSC-derived motor neurons did not cause noticeable changes in cell morphology or death. While the impact of BET inhibition on TAF15 needs to be determined, it and 3 additional members of the FUS complex mapped between the BRD4 and JQ1 regions of the ratio plot shown in Fig. 2c, suggesting proteostatic modulation by BRD4 and JQ1.

### BET inhibitor induced alterations to protein homeostasis of the BAF complex

The 2 megadalton mammalian BAF complex consists of up to 15 subunits which assemble combinatorially and in a tissue-specific manner to form distinct assemblies. Individual complexes differ in their sites of action in the genome and interactions with transcription factors or DNA repair/replication proteins [Alfert *et al*., 2019; Andrades *et al*., 2023]. Mutations in BAF subunits or alterations to their subunit compositions may cause lymphoid malignancies and other cancers [Mittal and Roberts, 2020; Shu *et al.,* 2020]. Several subunits of the esBAF complex likely interact with BRD4, but details about the role of BRD4 in BAF complex functions are sketchy and the influence of BET inhibitors on such complexes even less so.

Ten proteins belonging to the BAF family were among the UBQint proteins, 5 of which can interact with BRD4 (Fig. 5). ACTB was the most highly enriched in all samples (Table IV), 7 other BAF subunits were specific to the JQ1-AZD subsample, suggesting complex regulation at the protein level. With reference to the model presented in Fig. 2c, differential modulation of BAF proteins by BET inhibitors or BRD4 *via* the UPS is suggested (Table IV, column 2) - AZD5153 (3 proteins), JQ1 (1 protein), BRD4 (1 protein) or combinedly with BRD4 and JQ1 (1 protein). ARID1B is abrogated in JQ1 or AZD5153 treated samples, so either UPS modulation or ARID1B association with BRD-N *in vitro* is altered, suggesting multiple mechanisms. SMARCE1 was responsive to AZD5153 alone. In addition multiple histones, histone chaperones, and nucleosomal proteins were present in UBQint samples (not shown), which recruit the BAF complex to chromatin. The reasons why the esBAF subunits BRD9, BRG1, BCL11, BRD9, PBRM1, SMARCB1, SMARCD1-3, and ACT6LA were absent from the UBQint samples are not clear, but some preliminary conclusions were drawn from literature.

**Fig. 5.**
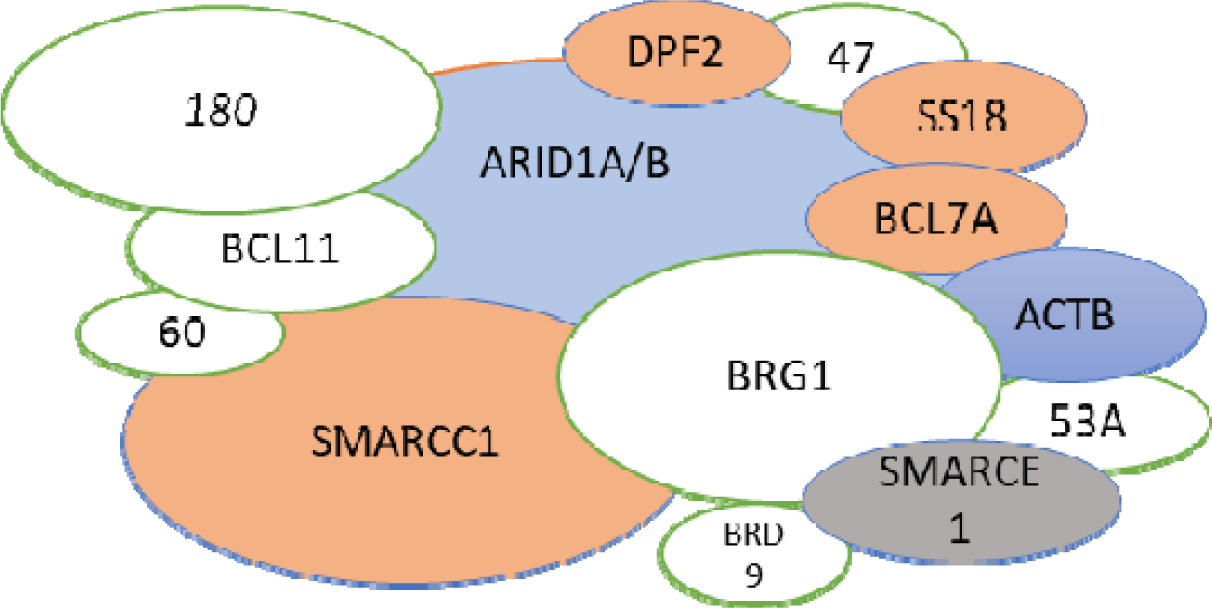
Representations of proteins known to function in the BAF complex identified in UBQint samples. Blue ovals – identified commonly in all samples, with thecaveat that ARI D1A can switch for ARID1B). Red ovals – Proteins commonly identified in JQ1 and AZD5153 treated samples. Grey oval – protein specific to AZD5153 treatment.

**Table IV.** Predicted responses of the identified BAF proteins to BET inhibitor or BRD4. Column 2 shows the proteins which may be responsive based on the model in Fig. 2c. Columns 3 thorough 6 show the number of optimized peptides identified for each protein in column 1, except ACTB, for which the total number of peptides are reported.

| BAF subunit | BET inhibitor influence zone | DMSO | JQ1 | AZD | I-BET |
| --- | --- | --- | --- | --- | --- |
| ACTB | n/a | 28 | 21 | 20 | 14 |
| ARID1A | BRD4 | 0 | 1 | 1 | 0 |
| ARID1B | JQ1; AZD | 2 | 0 | 0 | 1 |
| SS18 | AZD | 0 | 1 | 3 | 0 |
| DPF2 | AZD | 0 | 1 | 3 | 0 |
| BCL7A | JQ1-BRD4 | 0 | 2 | 1 | 0 |
| SMARCC1 | BRD4 | 0 | 1 | 1 | 0 |
| SMARCA2 | AZD | 0 | 1 | 3 | 0 |
| SMARCAD1 | JQ1 | 0 | 1 | 1 | 0 |
| SMARCE1 | n/a | 0 | 0 | 2 | 0 |

**Table V.** Shows predicted Myc responsive protein candidates with maximal changes across the BET inhibitor conditions. Column 2 indicates the known function from literature and column 3 the predicted response factor for each candidate protein.

| **UBQint candidate** | **Biological function** | **Responsive to** | **Reference** |
| --- | --- | --- | --- |
| Myc-R2 | Replication stress;<br>DNA damage | AZD5153 | Lam et al (2020) |
| Myc-R13 | Synaptic signaling | AZD5153; JQ1 | None |
| Myc-R3 | Nucleolar phosphoprotein | AZD5153; BRD4 | None |
| Myc-R5 | Multifunctional protein | BRD4 | None |
| Myc-R8 | Chaperone | BRD4 | None |
| Myc-R9 | Chaperone | BRD4 | None |
| Myc-R11 | Inflammation, chemotaxis,<br>apoptosis | BRD4 | None |
| Myc-R17 | Nucleosome structure | BRD4 | None |
| Myc-R20 | Antioxidant | BRD4 | None |
| Myc-R23 | Protease inhibitor | BRD4 | Zhang & Roeder |
| Myc-R32 | Epigenetic repression | BRD4 | None |
| Myc-R33 | Protein kinase regulator | BRD4 | None |
| Myc-R36 | Nucleosome structure | BRD4 | None |
| Myc-R14 | Deubiquitination | BRD4, JQ1 &<br>AZD5153 | None |
| Myc-R6 | Apoptosis | JQ1 | None |
| Myc-R10 | Chaperone | JQ1 | None |

Mashtalir *et al*. [2020] presented a three-dimensional (3D) structural model of the endogenously purified human canonical BAF complex bound to the nucleosome using a combination of single-particle cryo-EM, cross-linking mass-spectrometry (CX-MS), and homology modeling. They found that SMARCB1 in the “core module” abuts the nucleosome and connects to the “ARP module” via SMARCA4, both of which proteins were absent from UBQint samples. SMARCA2, a member of the catalytic “ATPase module”, interacts with the face of the nucleosome opposite that contacted by the core module. This module also consists of ACTB, which was the most abundant BAF subunit in UBQint samples, and ARID1A *via* its ARM 5-7 repeats (amino acids 1939–2282). While ARID1A and ARID1B can switch between each other in BAF complexes in cells, they appear to be mutually exclusive in BETi samples, with ARID1A detected in JQ1 and AZD5153 samples, and ARID1B only in DMSO and I-BET151 treated sample.

The BAF core module also includes SMARCC1/2, SMARCAD1, and SMARCE1; deletion of the former 2 subunits completely abrogates BAF assembly and functions [Mashtalir *et al*.]. The armadillo repeats in the ARID1A C-terminal domain [mapped by Sankaran *et al*., 2018] forms an interface with SMARCAD1 (CC and SWIB domains) and interacts with the SWIRM domain of SMARCC1, scaffolding the subunit arrangement into a core module of high rigidity. SS18 located in the core module connects with SMARCA4, providing structural context for the complex. DPF2, another protein in the core BAF complex and containing a plant homeobox domain, makes extensive contacts with SMARCB1, SMARCC1, and ARID1A/B, but can only associate with the fully assembled BAF core module [Mashtalir*et al*.]. BCL7A was suggested as a ‘bridge’ protein connecting the core complex with SMARCA4, but of unknown function.

The above information suggest that only the ‘core’ BAF complex was enriched in UBQint samples and that the partial complexes associate *in vitro* with BRD4-N. Extension of these results with further experimentation can help more clearly define the roles of BET inhibitors and BRD4 in the formation and regulation of BAF complexes, identify the E3 ligase(s) involved in modifying BAF subunits, and map the E3 ligase contact sites and the effects of mutations in BAF subunits. The findings also reiterate the utility of UBQuest for deeply probing proximity events and protein complexes in the ubiquitome, and help to purify the proteins to elucidate protein contacts. Profiling these proteins with additional BET inhibitors will help identify novel entry points and targets for modulating BAF complexes for therapeutic benefit. Since many BAF proteins are mutated in cancer and neurodegenerative diseases, densely mapping this space at the molecular, structural, regulatory, and pharmacological levels can provide large benefits.

### BET inhibitor mediated proteostasis alterations at the Myc promoter

The human Far Upstream Element (FUSE) Binding Protein 1 (FUBP1) is a multifunctional DNA and RNA binding protein and a master regulator of transcription, translation, and RNA splicing. FUBP1 hyperactivates Myc transcription by binding to the FUSE DNA element 1.5 kb upstream of the P1 promoter of Myc [Debaize and Troadec, 2019]. Of the many factors influencing Myc expression levels, lost binding of FUBP1 at the FUSE element has the highest correlation with Myc downregulation and the induction of differentiation in cells [Zaytseva & Quinn]. As a potent pro-proliferative and anti-apoptotic factor modulating complex networks, an oncoprotein, a tumor suppressor, and a protein with essential functions in hematopoietic stem cell maintenance and survival it is not surprising that dysregulation of FUBP1 expression leads to hematological disorders, solid tumors and neurodegenerative disorders. Gliomas, the most frequent cancers of the nervous system, frequently involve the combined deletions and monoallelic loss of FUBP1 and CIC, a transcriptional repressor, but the nature of interactions between the two proteins are not clear.

Pharmacological targeting of FUBP1 is an important objective in many diseases for regulating Myc expression [Dobrovolskaite *et al.,* 2022]. Other than that, FUBP1 also regulates expression of many additional genes [reviewed in Debaize & Troadec, 2019], notably RUNX1, SMN1 and TPT1, by binding to their promoters and to exon7 of SMN2 RNA [modulated by analogs of risidplam] *via* KH-elements in its protein structure [Meyer *et al*., 2021]. Using CRISPR screens, Elman *et al*. (2019) found that FUBP1 cooperates with other tumor suppressor genes to alter splicing and regulating N6-methyladenosine (m6A) RNA methylation, likely by physically associating with METTL3 [Elman *et al.,* 2019]. FUBP1 also represses translation of NPM1, PKD2, and GAP43 by binding to their 3’-untranslated regions [Debaize and Troadec].

There are no literature citations suggesting interaction between FUBP1 and BRD4. However, a disproportionately large signature of FUBP1 peptides among the mass spectrometry data (Fig. 6) suggests at least a networking effect between the two proteins. Preliminary analysis of the peptides suggests that JQ1 and AZD5153 may affect FUBP1 proteostasis in similar ways, likely involving UPS proteins. The compound specific protein interactions with FUBP1 appear to be quantitative in view of the number of FUBP1 peptides instead of stoichiometric due to recruitment into protein complexes, as may be the case with many other proteins in this study. A few other proteins interacting with FUBP1 were also among the UBQint proteins, some represented by large numbers of peptides and modulation with BET inhibition.

**Fig. 6.**
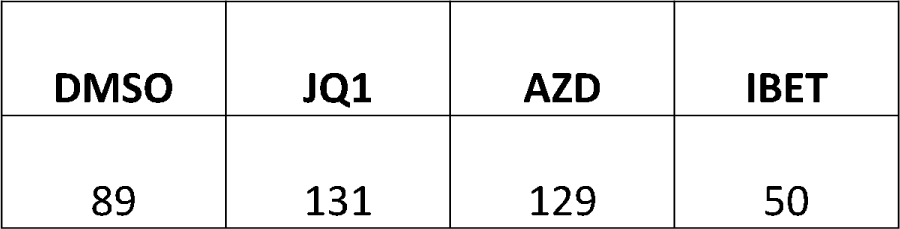
The table shows the total number of FUBP1 peptides identified with UBQint under each BET inhibitor treatment condition.

Identifying the E3 ligase activated by JQ1 and AZD5153 and mapping the site of its action on FUBP1 can help understand the effect of BET inhibitors on FUBP1 turnover. Further analysis of the UBQint samples can help understand if protein modifications alter the stability of FUBP1 or interactions with other proteins and RNA in cells. Indeed, FUBP1 is known to be subject to post-translational modifications such as, phosphorylation, acetylation, mono-methylation, di-methylation, sumoylation and ubiquitination, yet the roles of the modified forms of FUBP1 are poorly understood. A detailed understanding of the proteostasis mechanisms of FUBP1 and its modulation by BET inhibitors or BRD4 may suggest novel ways of therapeutically targeting Myc and other key regulatory proteins in lymphoid malignancies.

### Myc responsive gene products in UBQint samples suggest pleiotropic modulation of Myc gene expression by BET inhibitors

To identify Myc-responsive gene products and their modulation by BET inhibitors, UBQint proteins were queried for presence of such proteins by reference with the Gene Set Enrichment Analysis database [GSEA; Zeller *et al.,* 2003]. 52 UBQint proteins matched with the GSEA database entries suggesting alterations to Myc activity in the UBQint samples (Fig. 7), 11 of which were specific to the JQ1-AZD subsample. Sixteen proteins exhibited maximal induction based on peptide counts and comparisons with DMSO or I-BET samples, Table IV. Ten of these proteins are suggested to be modulated by BRD4, with reference to the ratio plot shown in Fig. 2c., and 2 additional proteins with contributions from at least one of JQ1 and AZD5153. On the same basis, 4 other proteins are suggested as being modulated by Myc directly by BET inhibitors. If so, the effects of BET inhibitor treated cells may be upstream or independent of BRD4 protein status.

**Fig. 7.**
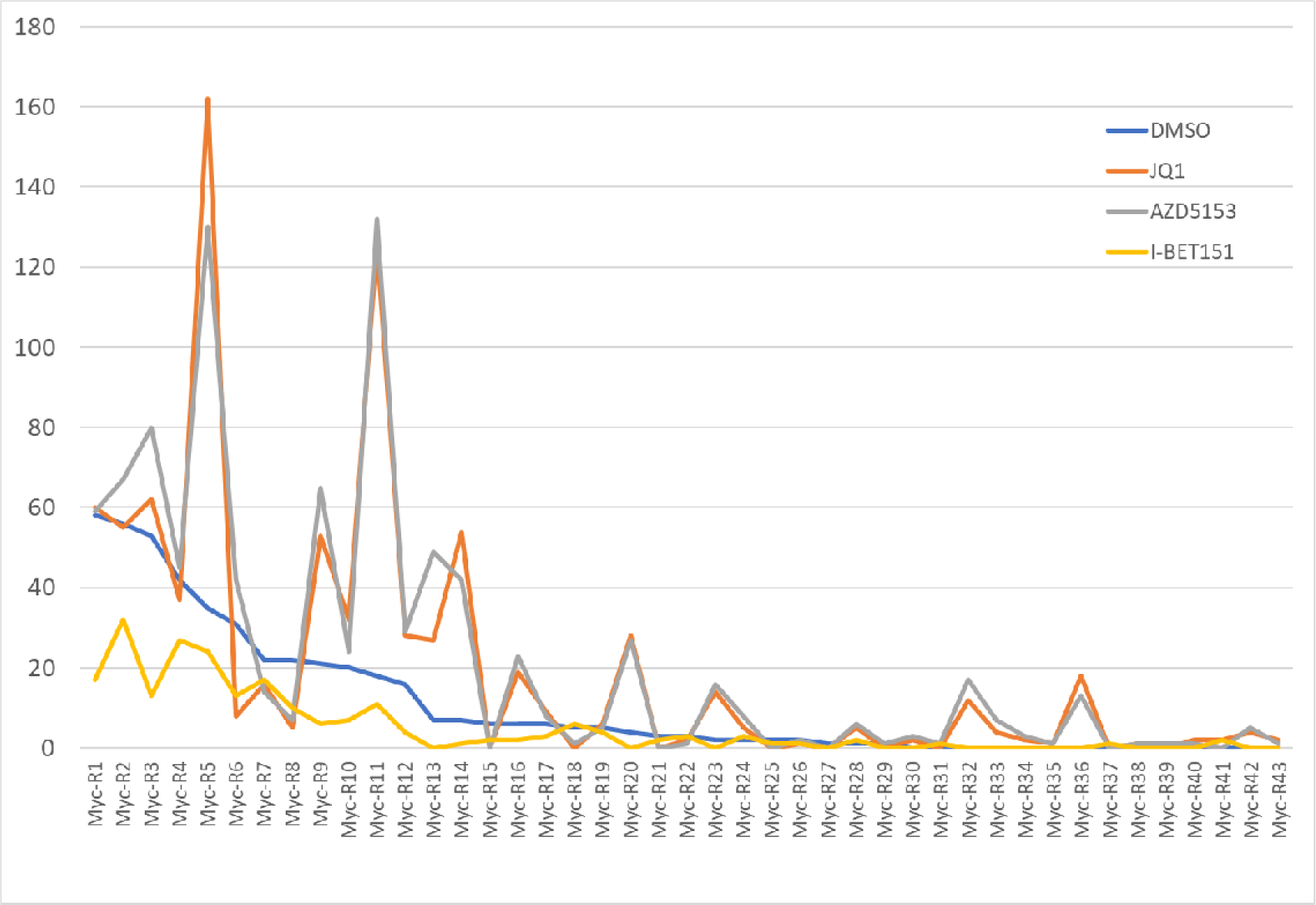
The chart shows total number of peptides identified for Myc responsive proteins. The X-axis indicates protein candidates; Y-axis shows the total number of peptides for each protein in the X-axis.

### Apoptotic proteins affecting BCL2 expression identified in UBQint samples

JQ1 has been shown to be a potent inducer of p53-independent apoptosis in the absence of its effects on Myc gene expression [Hogg *et al*., 2016]. The authors showed that JQ1 potently induced proteolytic cleavage of PARP1 as a downstream marker of caspase activation and overexpression of Bcl-2 in mice inhibited apoptosis induction by JQ1. They concluded that epigenetic regulation of endogenously expressed Bcl-2 family proteins mediated the apoptotic response to BET-inhibition, principally due to a shift in the ratio of Bim to Bcl-2 and Bcl-XL transcription. While that may be so, it still leaves mechanistic questions open, particularly for densely mapping the proteomic events (and protein contacts) leading from JQ1 treatment to apoptosis induction.

While a substantial signature of PARP1 degradation was observed in UBQint proteins, Bcl-2 or Bim were absent, suggesting the existence of alternate mechanisms. Mass spectrometry information suggests sensitivity of PARP1 to different E3 ligase(s), likely induced by JQ1 or AZD5153 as compared with the DMSO sample or I-BET151. A few other Bcl family proteins which recruit caspases, caspases, BH3-domain proteins, and caspase regulating proteins were also identified. In most cases differential sensitivity of these proteins towards JQ1 and AZD5153 were observed, suggesting alterations to UPS activity. The 29 pro- and anti-apoptotic proteins that were identified included many proteins acting independently of p53 or Bcl-xl family proteins, showing differential regulation in most cases by JQ1 or AZD5153. The significance of these findings and their contributions to BET inhibitor-induced apoptosis or acquired resistance to BET inhibition need to be elucidated.

### BET inhibitors exhibit a pronounced UPS triggering effect, however many additional compounds and FDA approved drugs may also exhibit the same property

To explore if the UPS triggering effect is a property shared by other small molecules, 14 additional compounds were tested in HEK293T cells. UBQuest proteins from the treated cells were subjected to a downstream ELISA assay, termed ubSCREEN, which utilizes subtractive proteomics to measure the relative abundance of UBQuest products from compound treated cells. The signal intensity greater than with control compounds or assay background indicates the presence of UPS proteins and/or their substrates; the same assay also captures the protein-UPS component responsible for the increased signal intensity.

As shown in Fig. 8, HEK293T cells treated with many compounds (represented by blue bars), including lenalidomide, showed high signal intensities in the ubSCREEN assay. This is expected with lenalidomide since it acts by modulating the substrate specificity of the CRL4C^RBN^ E3 ubiquitin ligase and induces degradation of many proteins, including IKZF1 and IKZF3 by CRL4^CRBN^ [Fink and Ebert, 2015]. The proteins giving rise to the signals have not been analyzed by mass spectrometry. Similar results were obtained with pomalidomide and iberomide (data not shown). BRD4-1, BRD4-2, and BRD4-3 compounds shown in Fig. 8 are JQ1, AZD5153, and I-BET151, respectively.

**Fig. 8.**
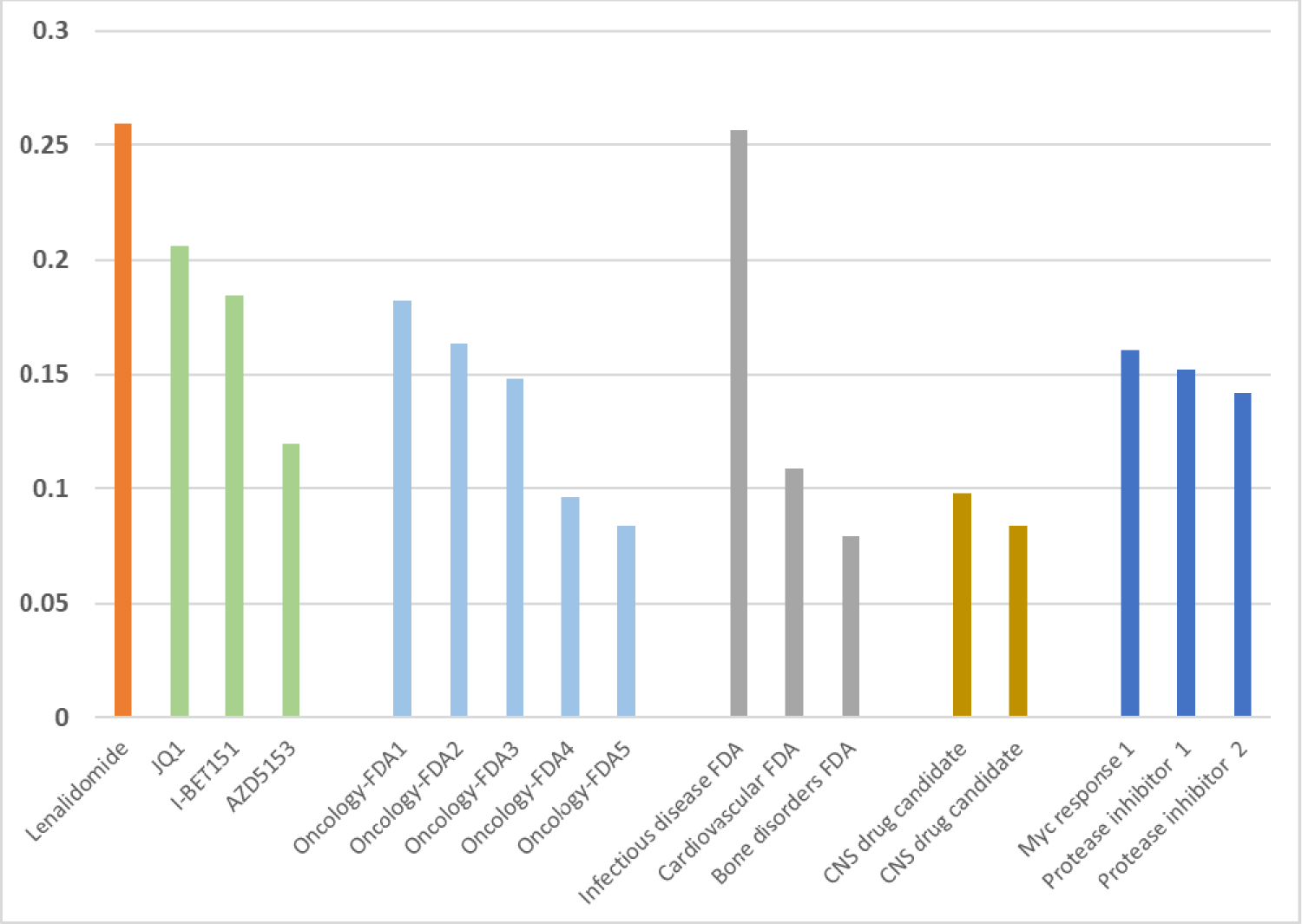
The chart shows ELISA signal intensities of the compounds and FDA approved drugs (X-axis) relative to a control sample (HEK293T cells treated with DMSO). A ratio cutoff of 0.1 was arbitrarily used to distinguish compounds which may trigger UPS effects, since BRD4-3 (I-BET151) has been confirmed in this study to exhibit a moderate effect, and the responsive E3 ligases and candidate substrates identified.

Interestingly, 4 additional FDA approved compounds exhibited signal intensities appreciably above background, whereas 4 other FDA approved compounds did not, indicating, 1) that some drugs may be effective in the clinic despite triggering the UPS, and 2) that the efficacy of these drugs and side effect profiles can perhaps be improved by identifying the proteins involved, and using combination therapies to downregulate them. Compound 1 was extensively tested for clinical use but eventually abandoned due to severe toxicity. It follows that UBQuest may be a useful tool for detecting compound-induced UPS triggers early during the drug discovery process to avoid costly failures later in the clinic and likely also to try to optimize the compounds via medicinal chemistry during early discovery phases to one that may retain efficacy but improved toxicity profile, particularly as it relates to UPS triggering effects for applications other than oncology therapeutics. The compounds and FDA drugs represented by grey bars in Fig. 8 likely do not alter the ubiquitome in HEK293T cells. It is not clear if higher doses of these compounds may trigger UPS effects. The ubSCREEN assay has been optimized in 96-format and is automatable.

## DISCUSSION

This is the first report describing a high throughput process for co-enrichment of proteostasis substrates and the UPS components involved. The proteins can be identified directly with mass spectrometry. However, since it was implemented in comparative mode with a cell line under differing physiological states, and since BET inhibitors were used as a means to alter the states, an additional step of selecting the proteins by physical association with a BRD4 fragment was employed. A surprising finding is that the BET inhibitors trigger UPS factors, giving rise to modified proteins. The second surprising aspect is that the modified proteins associate with BRD4 and many of the associations are compound-specific. Partial characterization of UBQint proteins identified many proteins known from literature which are involved in transcription and chromatin maintenance and complex with BRD4. The proteins co-purified several E3 ligases and accessory UPS proteins suggests, thereby implicating form and function. Extension of these findings over a time-course, in a range of cell types, and with sub-cellular fractionation, may reveal if functional links exist between BRD4-containing complexes and UPS mechanisms. Mono-ubiquitinated, SUMOylated, ufmylated, and ubiquitin binding proteins are believed to alter protein localization and interactions, so observing the protein modifications in selected candidates in a drug discovery setting can help understand their contributions, if any, to BRD4 function.

Whether the heightened UPS activity observed in presence of BET inhibitors is a compound-related effect or downstream of BRD4 action (BRD4-mediated) remains to be determined. The model presented in this report is based on differences in peptide count and helps to select some proteins for further analysis. These proteins can be profiled with ELISA and additional proteins involved in the complexes can be identified.

Susceptibility to new E3 ligases in response to BET inhibitors was a frequently observed theme in these studies. For instance, FUBP1 degradation in resting cells is consistent with suppressing Myc enhancer activity and an objective of BRD4 inhibition. However, the increased UPS activity towards FUBP1 under BET inhibited conditions suggests targeting by a new E3 ligase. Likewise, several proteins involved in Myc promoter activity and BAF subunit regulation appear to be subject to compound-induced UPS changes. The existence of these mechanisms under cellular conditions and elucidating the mechanisms involved are needed. Additionally, in view of the large UPS footprint observed in this study, repeating it at earlier time points can help segregate them into early, mid, or late responses. In turn, the observations can help determine the mechanistics involved and identify the key proteins responsible for the overall phenotypic effects of BET inhibition.

The co-purification of several protein kinases and phosphates in UBQint samples with varying BET inhibitor sensitivities suggests their involvement in BRD4 function. It is not clear to what extent the kinase activity of BRD4 itself is responsible for the proteomic changes observed. If phosphorylation and ubiquitination of the same candidates can be observed in this setting, it may resolve the question of whether phosphorylation acts as a trigger for UPS modification. If so, it can be determined with cell biology if the events are proximity-dependent requiring co-localization of the substrate and E3 ligase on chromatin, and additional proteins can be identified. The proteins mentioned in this report may serve as biomarkers of proteostasis useful for profiling in drug discovery samples.

The process described in this report can also be used to probe the structure function relationships of BRD4, as influenced by BET inhibitors. For instance, BRD4 activates homeobox genes at the transcriptional level, so some UBQint proteins may likely be HOX-response proteins. Indeed, several homeobox transcription factors were identified in this study which were also differentially responsive to BET inhibitors. Likewise, proteins involved in cellular responses to the environment, stress response factors, P body proteins, proteins involved in phase separations, and protein network alterations due to the cells cycling may account for some of the observed proteomic changes.

UBQint process is a sensitive tool for *de novo* identification of UPS activity by globally scanning the proteome and selectively enriching the active ubiquitome. It is ideally suited for observing time-sensitive biological processes wherein compartment-specific changes of protein structure or stability over small windows of time and space can trigger potent downstream events in absence of observable protein modification from steady-state considerations. Besides compound treatments, the process is also highly useful for detecting aberrations in biological samples involving dysregulated or mutated proteins, such as in some neurodegenerative diseases, including Parkinson’s disease. Protein homeostasis dysregulation is a factor in many additional diseases. The information gathered from UBQint can spur the creation of event-based, mechanistic drugs acting in appropriate cellular compartments as an alternative to quantitatively disrupting the cellular pools of proteins in non-target locations.

The insights derived from this study can be carried forward for finding the optimal target, small molecule and drug combinations with BET inhibitors. While some UBQint proteins are obvious candidates, additional proteins can be identified and validated under cellular conditions. The candidates may serve as standalone, or combination targets, and also spur the creation of novel assays for profiling proteins from responsive and non-responsive cell types along with RNAseq to study the early Myc responses. Where a specific compound is not available, it may be worthwhile applying the PINTAC process with select candidates [Nallur, 2022] to selectively knock out single proteins in surrogate assays to benchmark with BET inhibitor outcomes. PINTAC assays generate biomarker signatures of drug action which can be used for screening or optimizing compounds phenotypically.

## MATERIALS & METHODS

Cells and reagents: DOHH2 cells were purchased from Creative Biomart. Cells were grown at 5% CO2 in 90% RPMI 1640 medium containing 10% heat-inactivated fetal bovine serum (FBS) and Penicillin-Streptomycin (Pen-Strep) according to manufacturers’ recommended protocols. HEK293T cells were purchased from ATCC (Catalog no. CRL-3216). Where appropriate, cell proliferation was measured using CellTiter 96 AQueous One Solution Cell Proliferation Assay (G3582, Promega) according to the manufacturer’s protocol. Cell Lysis Buffer (10X) was purchased from Cell Signaling Technologies (Catalog no. #9803). Recombinant BRD4 N-terminal protein was purchased from GenScript (BRD4-N (49-460aa), His, Human; Catalog No. Z03187). Pierce antibody biotinylation Kit (90407) was purchased from ThermoFisher. Nitrocellulose membrane was purchased from Cytiva (Catalog no. 32HK60).

Compounds and drugs: The BET inhibitors, JQ1 (Catalog No.S7110), AZD5153 (Catalog No.S8344) and I-BET151 (GSK1210151A) (Catalog No.S2780) were purchased from SelleckChem. Lenalidomide was purchased from Apexbio Tech (Catalog no. A4211). Other compounds and FDA approved drugs employed in this study were purchased from approved vendors and dissolved in solvents recommended by the manufacturers.

Drug treatments and protein preparation: DOHH2 cells were grown to 70-90% confluence overnight (cultured in media without FBS). 2 million cell were treated with a BET compound or DMSO at a final concentration of 1/10^th^ of the published IC^50^ for BRD4 for each compound – JQ1 – 7.7 nM; AZD5153 – 300 nM; and I-BET150 – 79 nM, and incubated for 72 hrs. After the incubation cells were collected by centrifugation, washed 3 times in cold PBS, lysed in 2 ml of cell lysis buffer, and incubated for 30 minutes at room temperature with occasional vortexing. Whole cell proteins were collected by centrifugation and stored at −20 degrees in aliquots.

For the ubSCREEN assay, HEK293T cells were grown to 90 per cent confluence in DMSO medium containing 10% FBS and Pen-Strep at 5% CO2 in T25 flasks. The cells were treated with compounds at 1/10^th^ of IC^50^ for each compound as published or indicated by the manufacturers. Cells were incubated with drug for 72 hours, and processed as indicated above for DOHH2 cells. No appreciable cytopathic effects were observed under these conditions. Whole cell extracts were stored in aliquots at −20 degrees C.

Selection of UBQint proteins: UBQuest is a proprietary process. UBQuest proteins are validated for their expression using western blots of key proteins including BRD4 and a set of BRD4 interacting and BET inhibitor responsive proteins and normalized against housekeeping genes (GAPDH). Where known the UBQuest proteins are matched with literature information of E3 ligases which target them, and also other UPS proteins with which they interact. Select proteins are confirmed by protein knock out with PINTAC peptides (Nallur, 2022). The targeting peptide may be directed towards the E3 ligase identified in UBQint proteins or to a candidate substrate. The PINTAC treated cells are subjected to RNAseq to identify any changes to transcribed genes.

An aliquot of whole cell proteins was subjected to the UBQuest process. Proteins enriched with UBQuest were detected with mass spectrometry (not shown). UBQint proteins were prepared by incubating an aliquot of UBQuest proteins with BRD4-N protein immobilized on nitrocellulose filters. 20 to 100 ug of BRD4-N was incubated with nitrocellulose filter circles and incubated with shaking at room temperature for 2 hrs. The filters were then washed twice with cold PBS, and blocked with 3% non-fat dry milk in PBS. Blocked filters were rinsed with cold PBS and incubated UBQuest proteins in PBS containing 0.1% Tween20 for 6 hrs. at room temperature with gentle shaking. After incubation, the filters were washed 3 times with cold PBS containing 0.1% Tween20, and twice with cold PBS. Filters were transferred to microcentrifuge tubes and bound proteins were eluted in 200 ul PBS by heating at 60 degrees C for 15 minutes with frequenting vortexing. UBQint proteins were collected by centrifugation.

Mass spectrometry and data analysis: Mass spectrometry of UBQint protein samples was performed at vendor sites. Mascot software was used to identify proteins from LC-MS/MS data against the Swiss-Prot human database using a 1% false positive discovery rate (FDR).

Proteins represented in the UBQint samples were identified by comparing the peptide sequences from mass spectrometry with BLAST (https://blast.ncbi.nlm.nih.gov/Blast.cgi), and their putative functions were annotated with reference to CORUM (https://mips.helmholtz-muenchen.de/corum/), GeneCards (https://www.genecards.org/Search/Keyword?queryString = HSP90B1). Candidate protein complexes amongst UBQint proteins or BRD4 were predicted by comparisons with BioGRID (https://thebiogrid.org/) databases. selected peptides from mass spectrometry were annotated for the presence of protein modifications with reference to PhosphositePlus database (https://www.phosphosite.org/homeAction.action). UBQint proteins were annotated for potential interactions with BRD4 or other proteins with reference to GeneCards database.

Construction of BET inhibitor response model with optimized peptide counts: Peptides with the same amino acid sequence for proteins commonly identified in JQ1 and AZD5153 treated samples were analyzed in the following manner. 1 peptide each from the JQ1 and AZD5153 was electronically subtracted and the process iterated until no peptides remained with the same sequence in the data set. The remaining peptides were designated as optimized peptides, which have unique amino acid sequence in only one of the samples or both. Ratios of these peptides (multiplied by 100; per cent ratio) for each protein in the JQ1 sample was plotted against the per cent ratio of the same protein in the AZD5153 sample. A per cent ratio greater than 250 (2.5-fold higher in JQ1 sample) was arbitrarily assigned as being responsive to JQ1, a per cent ratio of 0.4 or less (2.5-fold lower in JQ1 sample) as being responsive to AZD5153, and a ratio of 1 to be responsive to BRD4.

Proteins in the predicted BRD4 response zone were matched with protein interactions databases to identify putative interactions with BRD4 and also with one another. First level interactors of proteins with putative BRD4 interacting function in this set were identified in this manner.

Myc responsive proteins known in literature were collected from The Myc Target Gene Database (http://www.myccancergene.org/site/mycTargetDB.asp) and Gene Set Enrichment Analysis (GSEA - https://www.gsea-msigdb.org/gsea/msigdb/human/geneset/DANG_MYC_TARGETS_UP.html) databases. To measure the effects of BET inhibitors on candidate Myc response genes identified in UBQint proteins, the total peptide counts for each protein in the mass spectrometry data for the candidate Myc response gene product were plotted from each of the BET inhibited samples.

## Acknowledgements

The research embodied in this report was conducted by Girish Nallur. The manuscript was written by Girish Nallur.

The author discloses no conflicts of interest.

The author thanks Nick Meanwell, PhD, and Dolatrai “Dinesh” Vyas, PhD, for critical reading of the manuscript and offering helpful comments and suggestions.

## Notes

### Competing Interest Statement

The authors have declared no competing interest.

